# Interdependent regulation of stereotyped and stochastic photoreceptor fates in the fly eye

**DOI:** 10.1101/865717

**Authors:** Adam C. Miller, Elizabeth Urban, Eric L. Lyons, Tory G. Herman, Robert J. Johnston

## Abstract

Diversification of neuronal subtypes often requires stochastic gene regulatory mechanisms. How stochastically expressed transcription factors interact with other regulators in gene networks to specify cell fates is poorly understood. The random mosaic of color-detecting R7 photoreceptor subtypes in *Drosophila* is controlled by the stochastic on/off expression of the transcription factor Spineless (Ss). In Ss^ON^ R7s, Ss induces expression of Rhodopsin 4 (Rh4), whereas in Ss^OFF^ R7s, the absence of Ss allows expression of Rhodopsin 3 (Rh3). Here, we find that the transcription factor Runt, which is initially expressed in all R7s, activates expression of Spineless in a random subset of R7s. Later, as R7s develop, Ss negatively feeds back onto Runt to prevent repression of Rh4 and ensure proper fate specification. Together, stereotyped and stochastic regulatory inputs are integrated into feedforward and feedback mechanisms to control cell fate.

## Introduction

Nervous systems are extremely complex, with some organisms having thousands of different neuronal subtypes. Specification of neuronal subtypes often occurs through a simple linear logic in which cells choose general fate class, neuronal type, and, finally, subtype. This logic appears to apply to the human retina, where neurons decide between photoreceptor (PR) or non-PR fate classes. PRs then select either a motion-detecting rod or a color-detecting cone fate. Cones choose between blue and red/green subtype fates and, finally, red/green cones diversify into red or green fates. Often, transcription factors control these decisions; for example, CRX determines PR vs. non-PR fate, and NRL distinguishes rod vs. cone fate (Bessant et al., 1999; Furukawa et al., 1997; Mears et al., 2001; Rehemtulla et al., 1996; Viets et al., 2016). Many such decisions appear to be deterministic. However, sensory systems, in particular, also diversify cell fates by the use of stochastic cell fate decisions (Johnston and Desplan, 2010; Urban and Johnston, 2018). In the human retina, the final choice between the red and green cone fates appears to occur by chance. A noncoding regulatory DNA element called a Locus Control Region (LCR) is hypothesized to randomly loop to the promoter of either the red or green opsin gene, activating its expression and preventing expression of the other opsin (Smallwood et al., 2002a). Similar LCR elements are thought to control the stochastic selection of one of 1300 odorant receptors for expression in olfactory neurons in mice (Markenscoff-Papadimitriou et al., 2014; Monahan and Lomvardas, 2015; Serizawa et al., 2003). Stochastic cell fate specification is even observed in the olfactory system of the nematode *C. elegans*, whose development is highly stereotyped. Ca^2+^-mediated lateral inhibition randomly specifies fates of the two AWC olfactory neurons (Alqadah et al., 2016; Chuang et al., 2007; Troemel et al., 1999). Thus, stochastic mechanisms are widely utilized to diversify neuronal subtypes. We are interested in understanding how stochastic mechanisms are incorporated into gene regulatory networks to control cell fate.

Like PRs in the human eye, PRs in the fly eye are thought to be specified in a simple series of fate decisions (**Fig. 1A**)(Morante et al., 2007). PRs are differentiated into color-detecting inner PRs or motion-detecting outer PRs (Domingos et al., 2004; Mollereau et al., 2001). Inner PRs are further specified into R7 or R8 neuronal types (Cook et al., 2003; Xie et al., 2007). Finally, R7s and R8s are diversified into subtypes (Hsiao et al., 2013; Johnston and Desplan, 2014; Jukam and Desplan, 2011; Jukam et al., 2016; Jukam et al., 2013; Mikeladze-Dvali et al., 2005; Thanawala et al., 2013; Wernet et al., 2006; Yan et al., 2017).

**Figure 1.**
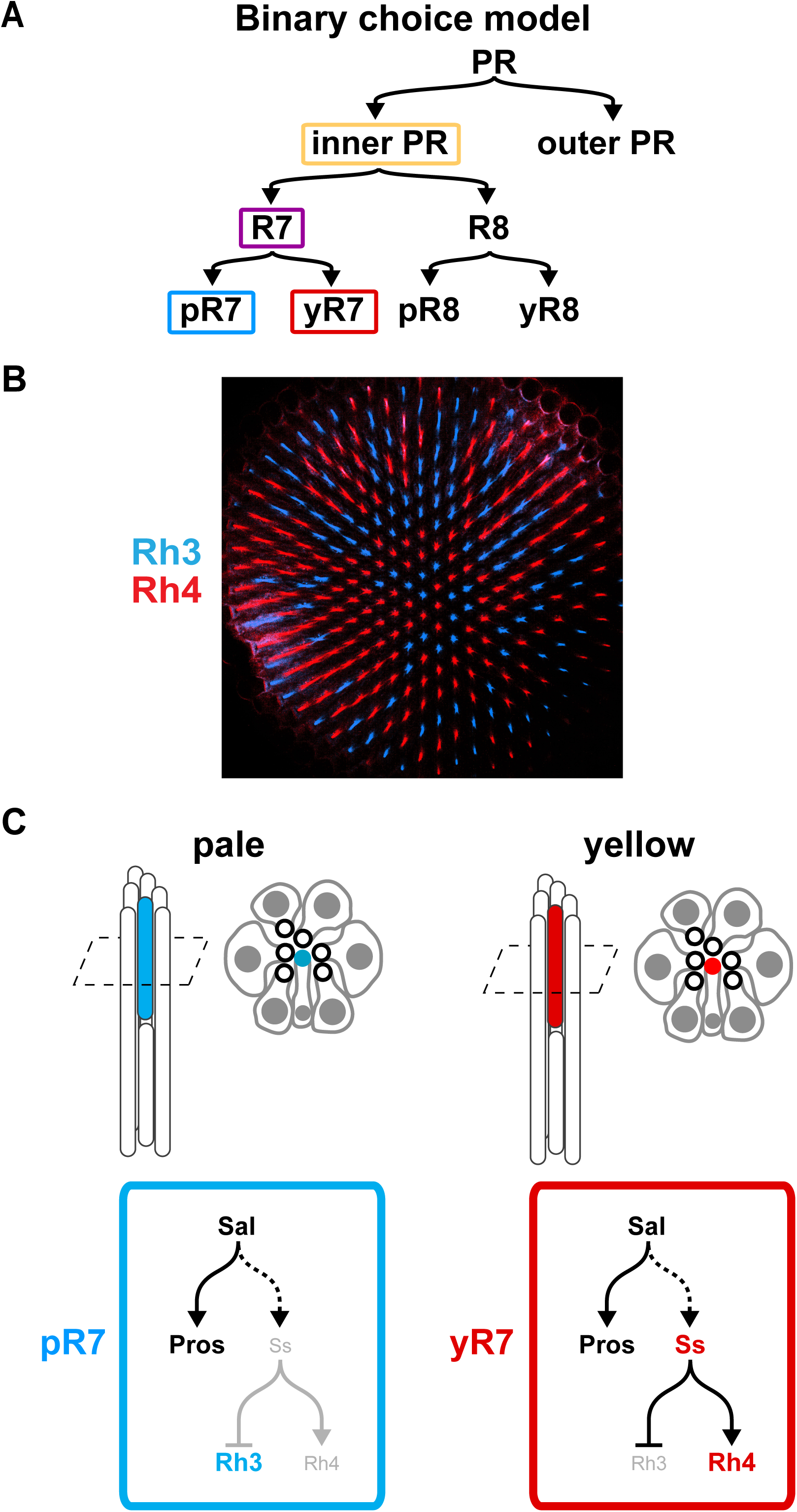
Photoreceptor subtype specification in *Drosophila melanogaster*. A. Canonical binary choice model for photoreceptor (PR) subtype specification. B. Stochastic pattern of Rh4-expressing yR7s and Rh3-expressing pR7s in a wild type adult fly retina. Rh4 (red), Rh3 (blue). C. Schematic and cross section of a single ommatidia containing R1-R8 cells. R7 cells denoted with Rh4 (red) or Rh3 (blue). Gene regulatory pathways responsible for pale vs. yellow R7 fate.

The R7 photoreceptor subtypes of the fly eye comprise a random mosaic (**Fig. 1B**)(Bell et al., 2007). This random distribution is controlled by the stochastic expression of the bHLH transcription factor Spineless (Ss). Ss is expressed in ~65% of R7s and induces yellow R7 (yR7) fate, including activation of Rhodopsin 4 (Rh4) and repression of Rhodopsin 3 (Rh3)(**Fig. 1C**). In the complementary ~35% of R7s where Ss is not expressed, R7s take on pale R7 (pR7) fate, marked by expression of Rh3 and absence of Rh4 (**Fig. 1C**) (Anderson et al., 2017; Johnston and Desplan, 2014; Wernet et al., 2006). The Spalt transcription factors (Sal) activates stereotyped expression of the general R7 fate gene Prospero (Pros) in all R7s and the stochastic expression of Ss (**Fig. 1C**). We here investigate how this stochastic regulatory mechanism is integrated into the gene regulatory network that specifies R7 fate.

Previous work has shown that the Run transcription factor is expressed in inner PRs (i.e. R7s and R8s) and induces aspects of inner PR fate, including the generation of small rhabdomeres and axonal targeting to the medulla (Kaminker et al., 2002). Ectopic expression of Run in outer PRs induces Rh3 reporter expression, but not Rh4 reporter expression (Edwards and Meinertzhagen, 2009), leading us to speculate that Run might participate with Ss in specifying R7 subtype.

Here, we describe the regulatory relationship between Ss and Run that yield PR subtypes in the fly eye and identify a complex gene regulatory logic that controls stochastic cell fate specification. We find that Run is sufficient to activate stochastic expression of Ss. Later, Run expression is restricted to pR7s lacking Ss. Negative feedback from Ss is necessary and sufficient to ensure proper Run expression during terminal differentiation. Perturbing this feedback by extending Run expression leads to misregulation of Rhodopsin expression, specifically the loss of Rh4 in yR7s. Additionally, we find that ectopic expression of the inner PR factors Run or Sal is sufficient to activate stochastic expression of Ss separate from general R7 fate. This finding suggests that inner PR regulators induce general and stochastic R7 fates features in parallel, challenging the established, simple model of binary decisions. Our studies reveal a complex interplay between stereotyped and stochastic regulators that controls PR specification in the fly eye.

## Results

### Ss and Run expression dynamics suggest regulatory interactions

We first examined the temporal dynamics of Ss and Run expression in 3^rd^ instar larvae. The developing fly eye is comprised of dividing undifferentiated cells. A morphogenetic furrow moves from posterior to anterior, followed by specification of PRs. Immediately following R7 recruitment, Run is expressed in all R7s, but Ss expression has not yet begun (**Fig. 2A, C-D**). Several rows later, Run expression persists in all R7s and Ss is expressed in ~65% of R7s (**Fig. 2A, C-D**)(Johnston and Desplan, 2014). At this stage, Ss and Run are co-expressed in yR7s, while only Run is expressed in pR7s. At 12 hours after puparium formation (APF), Run expression begins to turn off in Ss-expressing yR7s (**Fig. 2A, C-D**). Finally, by 48 hours APF, Ss and Run expression are mutually exclusive, leaving two classes of R7 cells: yR7s that express Ss only and pR7s that express Run only (**Fig. 2A-D**). Based on the expression dynamics of Run and Ss, we hypothesized that initially, Run activates stochastic expression of Ss and later, Ss feeds back to repress Run in yR7s (**Fig. 2D**).

**Figure 2.**
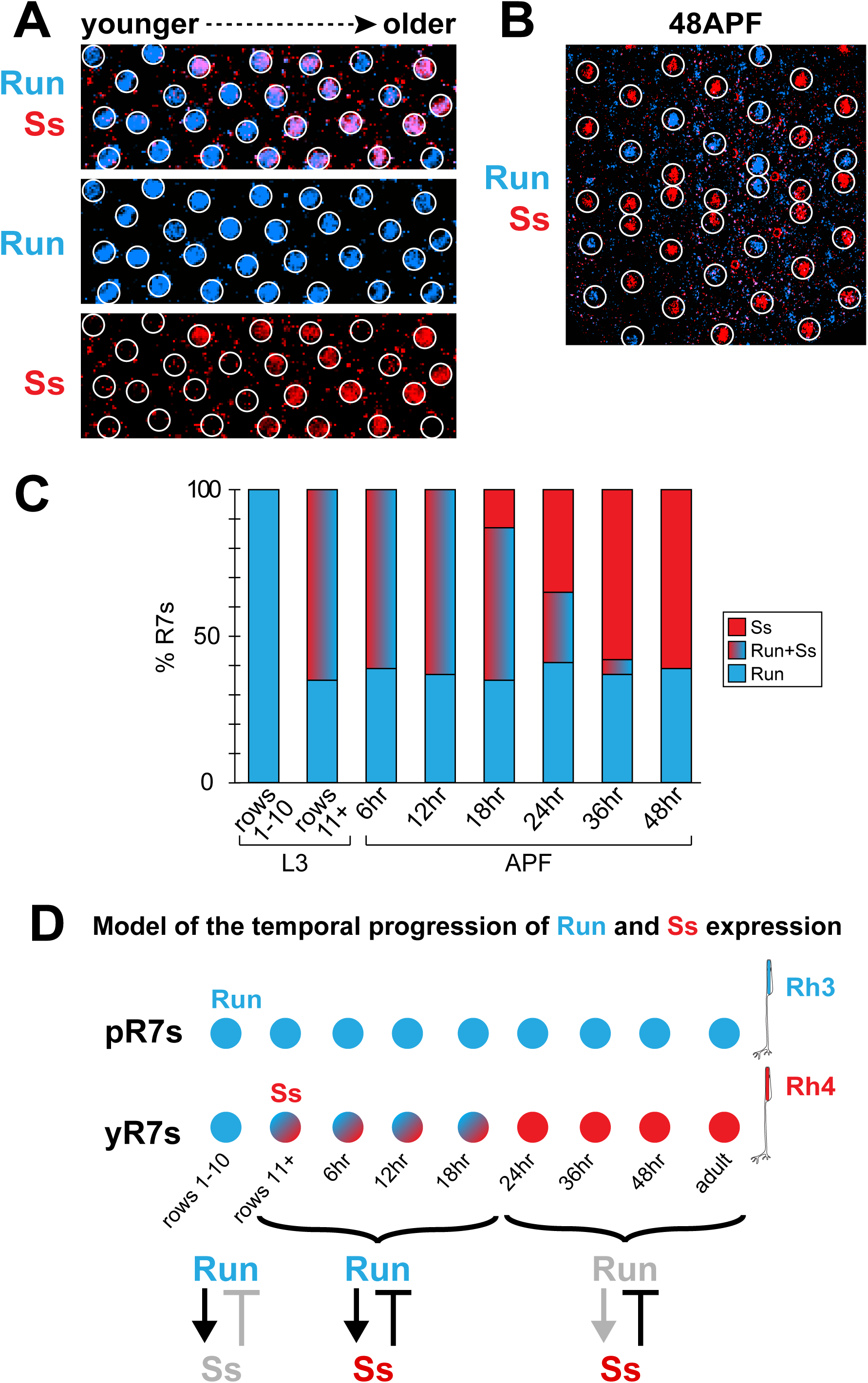
Run is restricted to Ss^OFF^ cells during development. A. Run (blue) is expressed in several rows of R7s before Ss (red) is expressed in a subset of R7s in the 3^rd^ instar larval retina. B. Run and Ss are exclusively expressed in subsets of R7s at 48 APF. C. Quantification of the temporal dynamics of Ss and Run expression. D. Model of the temporal dynamics of Ss and Run expression in pR7s and yR7s during development.

### Run activates R7 fate and stochastic expression of Ss in parallel

We next examined how Run regulates R7 fate features. Ectopic expression of Run in all PRs early in development caused severe morphological defects including a loss of the interommatidial space, precluding analysis at the single cell level (**Fig. 3A-B**). Nevertheless, we found that ectopic Run expression induces Rh3 in cells of all ommatidia and completely eliminates Rh4 expression (**Fig. 3B**). Thus, Run is initially expressed in all R7s, but it induces expression of the pR7 marker Rh3 while antagonizing expression of the yR7 marker Rh4.

**Figure 3.**
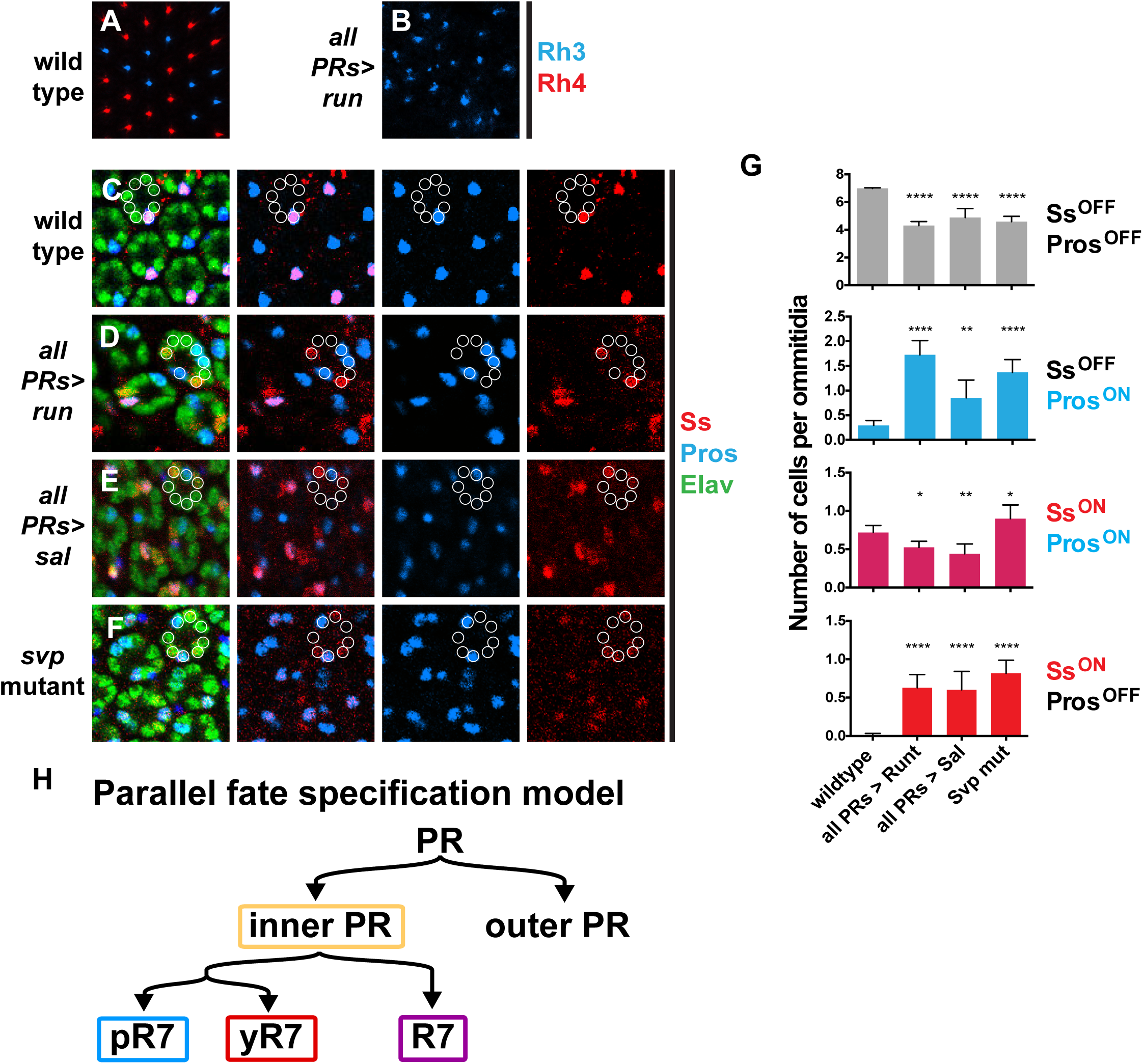
Run activates stochastic expression of Ss. A. Random distribution of Rh3 and Rh4-expressing R7s in the fly retina. B. Expression of Runt in all PRs induces Rh3 and represses Rh4. C-F. Ss (red); Elav (green) indicates PRs; Pros (blue) indicates general R7 fate; white circles denote each PR in an ommatidial cluster. C. Ss is expressed in a subset of R7s (marked with Pros) in wild type retinas. D-F. Flies with ectopic expression of Run, ectopic expression of Sal, or *svp* mutation displayed Ss and Pros in random subsets of PRs leading to cells with Ss^OFF^ Pros^OFF^, Ss^ON^ Pros^ON^, Ss^OFF^ Pros^ON^, or Ss^ON^ Pros^OFF^. D. Ectopic expression of Run. E. Ectopic expression of Sal. F. *svp* mutants. G. Quantification of F-I. H. Model of how the inner PR regulators Run and Sal control stochastic and general R7 features in parallel.

This observation suggested two possible relationships among Run, R7 fate, and Ss expression. Run could induce R7 fate and repress Ss, preventing expression of Rh4. Alternatively, Run could induce R7 fate, including stochastic expression of Ss, and then act downstream of Ss to repress Rh4. To distinguish between these possibilities, we ectopically expressed Run in all PRs and analyzed Ss expression. We observed expression of Ss in a random subset of PRs (**Fig. 3C-D, G**), suggesting that Run activates stochastic expression of Ss and then plays additional roles downstream of Ss to regulate Rh4 expression.

Ectopic expression of Run during early differentiation in larvae also induced Pros, an R7 fate marker, in a random subset of outer PRs (**Fig. 3C-D, G**), in contrast to expression in late pupal stages, which does not induce Pros (Kaminker et al., 2002). Interestingly, we saw all possible combinations of Ss and Pros expression, including Ss^OFF^ Pros^OFF^, Ss^OFF^ Pros^ON^, Ss^ON^ Pros^ON^, and Ss^ON^ Pros^OFF^ cells (**Fig. 3C-D, G**). We observed similar phenotypes in flies with ectopic expression of Sal (an inner PR transcription factor) and in *sevenup (svp)* mutants, in which Sal and inner PR fate are derepressed (**Fig. 3E-G**) (Miller et al., 2008; Mlodzik et al., 1990). In all three conditions, the number of Ss^OFF^ Pros^OFF^ cells per ommatidium decreased, consistent with a conversion of outer PR fate to a different fate (**Fig. 3G**). Ss^OFF^ Pros^ON^ cells increased, suggesting that general R7 fate is derepressed (**Fig. 3G**)(Miller et al., 2008; Mlodzik et al., 1990). The number of Ss^ON^ Pros^ON^ cells decreased upon ectopic expression of Run and Sal and increased in *svp* mutants, suggesting that complex regulatory interactions and/or genetic background altered the penetrance of Ss and Run expression (**Fig. 3G**). Finally, Ss^ON^ Pros^OFF^ cells increased (**Fig. 3G**), showing that inner PR regulators can activate stochastic Ss expression in the absence of general R7 features. Consistent with this observation, stochastic Ss expression is unaffected in *pros* mutants (Johnston et al., 2011). Together, these data suggest that inner PR regulators activate stochastic and general R7 features in parallel (**Fig. 3H**) in contrast to the binary choice model of PR specification (**Fig. 1A**).

To understand how Run regulates Ss expression, we assessed Run paralogs and interacting partners. Run is a BTB transcription factor whose paralog is Lozenge (Lz), a critical regulator of R7 fate (Daga et al., 1996). Run and Lz act with cofactors Brother (Bro) and Big brother (Bgb) to regulate target gene expression (Kaminker et al., 2001; Li and Gergen, 1999). We tested whether these factors are necessary for expression of Ss, Rh3, and Rh4. *run* null mutant clones and *run* RNAi display wild type Rh3 and Rh4 expression (**Fig. S1A-B, D, F**). *lz* null mutants display a near complete loss of R7s, as previously reported (Daga et al., 1996), yet Rh3 and Rh4 are stochastically expressed in the remaining cells (**Fig. S1C**). Further, loss-of-function *bro* mutants had no significant change in the ratio of Rh3- and Rh4-expressing R7s (**Fig. S1E-F**), suggesting that the family of BTB transcription factors have redundancy in their regulation of inner PR fate. Overexpression of Run, Bro, or Bgb specifically in R7s had no effect on the ratio of Ss^ON^ to Ss^OFF^ cells (**Fig. S1G**), suggesting that the levels of these regulators do not determine the ratio of stochastic Ss^ON/OFF^ expression. We conclude that Run is sufficient but not necessary to induce general R7 fate and stochastic *ss* expression.

### Ss restricts Run expression to Ss^OFF^ cells later in development

Our data suggested that Run activates Ss initially and Ss feeds back to repress Run later in development (**Fig 2D**). If Ss feeds back to repress Run, then Run should be derepressed in *ss* mutants. Indeed, in whole eye null mutant clones of *ss*, Run was expressed in all R7s (**Fig. 4A-B**). To test the cell autonomy of this effect, we examined *ss* null single-cell clones and observed nearly complete derepression of Run (**Fig. 4C, E**). We saw similar effects on Rhodopsin expression in adult R7 photoceptors as *ss* null single-cell clones displayed loss of Rh4 and gain of Rh3 expression (**Fig. 4F, H**). Thus, Ss is required in yR7s to repress Run.

**Figure 4.**
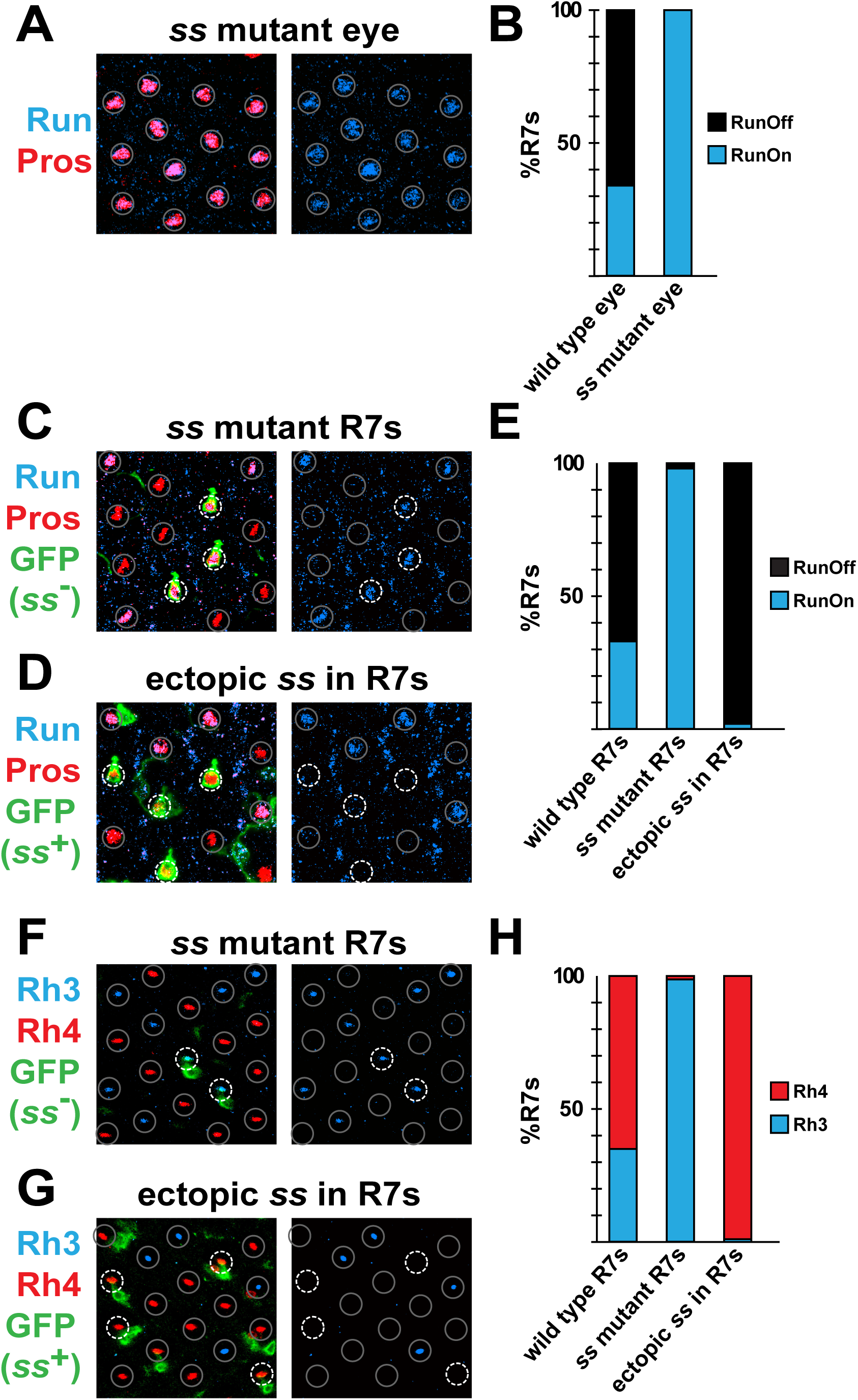
Ss restricts Runt to Ss^OFF^ cells. A. Run was expressed in all R7s in *ss* null mutant eyes. R7 cells marked by gray circles. B. Quantification of A. C. Run was de-repressed in *ss* null mutant R7s. Wild type R7 cells are indicated with gray circles. GFP (dashed white circles) marks single cell R7 *ss* mutant clones. Single cell clones also occur in non-R7 cells (GFP, no circle). D. Run was repressed in R7s with ectopic Ss expression. Wild type R7 cells are indicated with gray circles. GFP (dashed white circles) marks single cell ectopic expression of Ss in R7s. Single cell clones also occur in non-R7 cells (GFP, no circle). E. Quantification of C and D. F. Rh3 was de-repressed in *ss* null mutant R7s. Wild type R7 cells are indicated with gray circles. GFP (dashed white circles) marks single cell R7 *ss* mutant clones. Single cell clones also occur in non-R7 cells (GFP, no circle). G. Rh3 was repressed in R7s with ectopic Ss expression. Wild type R7 cells are indicated with gray circles. GFP (dashed white circles) marks single cell ectopic expression of Ss in R7s. Single cell clones also occur in non-R7 cells (GFP, no circle). H. Quantification of F and G.

As Ss is necessary and sufficient for yR7 fate (Johnston et al., 2011; Wernet et al., 2006), we predicted that Ss would be sufficient to repress Run in R7s. Indeed, ectopic expression of Ss in pR7s caused near-complete repression or Run (**Fig. 4D-E**). Likewise, ectopic expression of Ss in single R7s induced Rh4 expression and repressed Rh3 expression (**Fig. 4G-H**). Together, these data show that Ss represses Run in mature yR7s.

### Run is repressed in Ss^ON^ R7s to allow Rh4 expression

We next wondered whether repression of Run by Ss in yR7s is required for proper Rh3 and Rh4 expression. We hypothesized that ectopically expressing Run in adult yR7s might perturb Rh3 and/or Rh4 expression. Indeed, ectopic expression of Run in yR7s caused significant repression of Rh4, leading to “empty” R7s that expressed neither Rh3 nor Rh4 (**Fig. 5A-E**).

**Figure 5.**
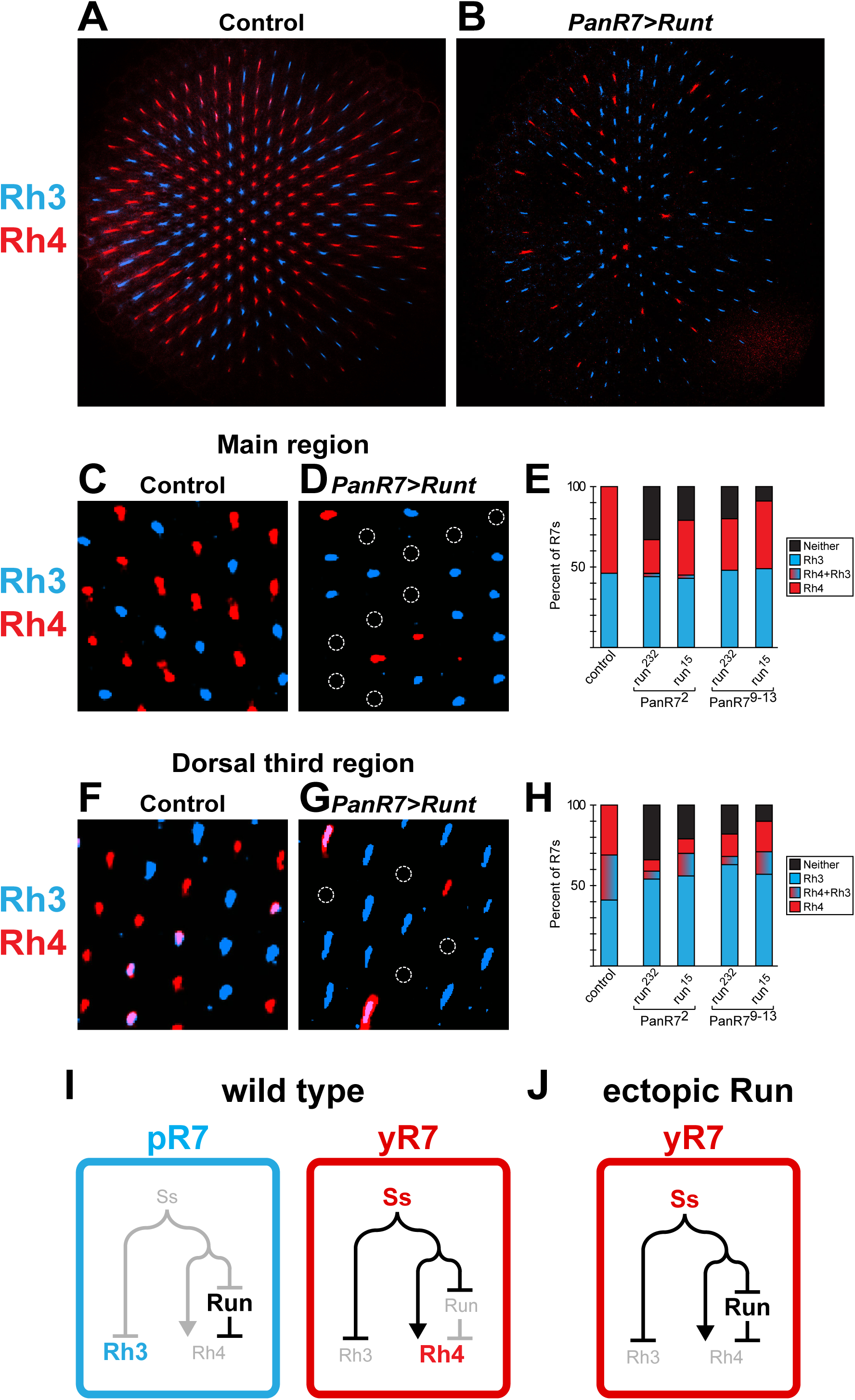
Runt is repressed in Ss^ON^ cells to allow Rh4 expression. A. Expression of Rh3 and Rh4 in a wild type retina B. Extending Run expression in yR7s represses Rh4. C. In the main region of the retina, Rh3 and Rh4 are exclusively expressed. D. Extending expression of Run represses Rh4, leaving “empty” R7s in the main region of the retina. E. Quantification of C-D. F. In the dorsal third region of the retina, Rh3 is co-expressed with Rh4 in yR7s. G. Extending expression of Run represses Rh4, generating Rh3-expressing “pseudo pR7s” in the dorsal third of the retina. H. Quantification of F-G. I. Model for the relationship between Ss, Run, Rh3, and Rh4 expression late in R7 subtype specification. J. Model for how extending Run expression in yR7s leads to Rh4 repression.

Expression of Rh3 and Rh4 is mutually exclusive throughout the majority of the eye (**Fig. 5A, C**). However, in the specialized dorsal third region, Rh3 is activated in Rh4-expressing yR7s (**Fig. 5F, H**)(Mazzoni et al., 2008). Ectopic Run expression led to a loss of Rh4 in dorsal third R7s, yielding “pseudo pR7s” that expressed Rh3 (**Fig. 5G-H**). Together, these data argue that Ss represses Run in yR7s to permit expression of Rh4 (**Fig. 5I-J**).

## Discussion

Our characterization of the regulatory relationship between the transcription factors Ss and Run revealed a surprising complexity in the logic controlling PR specification. Run activates stochastic expression of Ss, which feeds back to repress Run and prevent Rh4 repression. Moreover, we find that inner PR factors (Run and Sal) induce stochastic expression of Ss in parallel with general R7 fate, challenging the canonical binary choice model for PR specification in flies.

Our data support a temporally dynamic model of stochastic R7 subtype specification. Initially, Sal and Run are expressed in all R7s. They activate the expression of Ss in a random subset of R7s. In yR7s, Ss induces Defective Proventriculus (Dve), a transcriptional repressor that turns off Rh3 expression. Ss directly induces Rh4 and feeds back to repress Run, allowing derepression of Rh4. In pR7s lacking Ss, Sal represses Dve and feeds forward to activate Rh3 expression. In the absence of Ss, Run remains expressed (**Fig. 6**) (Johnston et al., 2011; Thanawala et al., 2013). This model supports a complex interplay between stereotyped factors like Run and stochastic factors like Ss.

**Figure 6.**
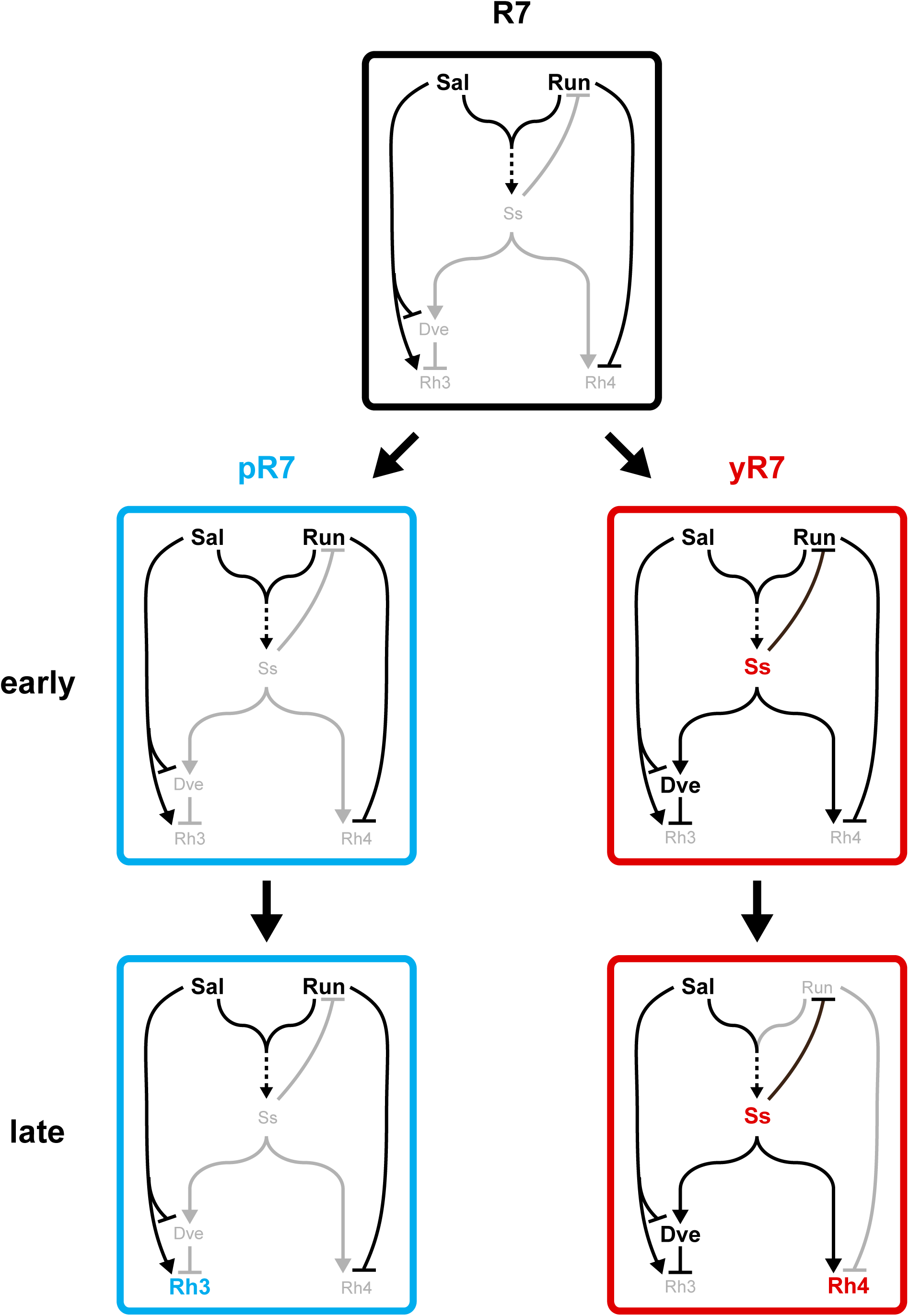
Model for temporal specification of R7 subtypes. Sal and Run activate inner cell fate and R7 fate. Sal and Run activate stochastic Ss expression early in development in parallel with R7 fate. Later in development, Ss interacts with Sal and Run in incoherent feed forward loops to specify pR7 vs. yR7 fate. Ss feeds back to repress Run in yR7s to allow Rh4 expression.

Though both Run and Sal induce stochastic Ss expression, they differ in their response to Ss itself. Sal is expressed in all R7s throughout development, indifferent to Ss expression, and is critical for Rh3 expression in pR7s (Johnston et al., 2011) and the generation of small rhabdomeres in all R7s (Mollereau et al., 2001). Run, on the other hand, is repressed by Ss in yR7s to allow Rh4 expression. The network is further complicated since Sal activates expression of Run (Domingos et al., 2004).

How Sal and Run genetically interact with Ss to regulate Rh3 and Rh4 differs in complex ways. Sal activates Ss to repress Rh3 (via Dve) and Sal also feeds forward to activate Rh3. Run activates Ss to induce Rh4 and also feeds forward to repress Rh4. Both Run and Sal interact with Ss in incoherent feedforward loops, suggesting that stereotyped and stochastic mechanisms are in direct competition for PR fate (**Fig. 6**).

Stochastic mechanisms are often hypothesized to be simply added onto or modified from existing gene regulatory mechanisms during evolution. For example, in human color PRs, the genes encoding the red and green opsins are different alleles of the same gene in ancestral primate species. It is thought that a recombination event brought these genes onto the same chromosome. A shared LCR putatively loops to the promoter of only one opsin gene per cell, leading to stochastic opsin expression (Jacobs and Nathans, 2009; Nathans, 1999; Smallwood et al., 2002b).

Our finding that Run activates Ss and then Ss feeds back to repress Run, suggests that this stochastic specification mechanism required significant regulatory network changes during evolution. Most likely, stochastic expression of Ss evolved first. There would be no effect on Rh4 or Rh3 expression since Runt would be expressed in all R7s to prevent Rh4 expression and Ss could not activate the repressor Defective Proventriculus (Dve) to inhibit Rh3. Either activation of Rh4 (requiring repression of Run) or activation of Dve to repress Rh3 would then evolve. Analysis of different insect species may identify the ancestral state and how these regulatory events evolved (Wernet et al., 2015). For example, if activation of Rh4 coupled with Run repression evolved first, we would expect to find species with expression of Rh3 alone in a subset of R7s and co-expression of Rh4 and Rh3 in a complementary subset of R7s. In contrast, if repression of Rh3 evolved first, we would see species with expression of Rh3 in a subset of R7s and no Rh expression in the complementary subset, which seems unlikely.

Run and Sal induce inner PR fate. Interestingly, they induce stereotyped R7 fate (i.e. Pros expression) and stochastic subtype fate (i.e. Ss expression) in parallel, independent pathways. Our findings conflict with the canonical view that PR specification is a simple decision tree from outer PR/inner PR class to R7/R8 type to yR7/pR7 subtype. Regulation of stochastic expression may provide flexibility to alter fates between PRs rapidly during evolution. Butterflies provide a beautiful example of this flexibility, where their two “R7s” in each ommatidium each make independent random fate choices based on Ss expression (Perry et al., 2016).

In conclusion, our studies identify a regulatory network controlling stochastic fate specification. These findings change our long-standing view of fate specification in the fly eye from a simple decision tree to a complex network of parallel fate choices. Our work makes specific predictions about the evolution of the eye, awaiting investigation through phylogenetic characterization of Rh expression patterns.

## Materials and Methods

### *Drosophila* genotypes and stocks

Flies were raised on standard cornmeal-molasses-agar medium and grown at 25°C.

| Short Genotype | Complete Genotype | Figures | Source |
| --- | --- | --- | --- |
| *wild type* | *yw*; +; + or +; +; + | 1B, 2A-C, 3A, 3C, 3G, 5A, 5C, 5E-F, 5H |  |
| *all PRs>run* | *yw*; *IGMR>Gal4/+*; *UAS>run/+* | 3B, 3D, 3G, 3G |  |
| *all PRs>sal* | *yw*, *UAS>sal*; *IGMR>Gal4/+*; + | 3E, 3G | (Kuhnlein and Schuh, 1996) |
| *svp* mutant | *ey>Flp*; +; *FRT82b svp<sup>2</sup>/FRT82b GMR>hid*, *cL* | 3F-G | (Stowers and Schwarz, 1999) |
| *ss* mutant eye | *ey>Flp*; +; *FRT82b ss<sup>d115.7</sup>/FRT82b GMR>hid*, *cL* | 4A-B | (Duncan et al., 1998; Stowers and Schwarz, 1999) |
| *ss* mutant R7s | *GMR>flp/w*; +/*Act>Gal4*, *UAS>mCD8-GFP*; *FRT82b ss<sup>d115.7</sup>/FRT82b*, *tub-Gal80* | 4C, 4E-F, 4H |  |
| ectopic *ss* in R7s | *GMR>flp/w*; *UAS>ss/Act>Gal4*, *UAS>mCD8-GFP*; *FRT82b/FRT82b*, *tub-Gal80* | 4D-E, 4G-H |  |
| *panR7>run* | *yw*; *panR7>Gal4/+*; *UAS>run/+* or *yw*; *panR7>Gal4/+*; *UAS>run/lroC>LacZ*<br>Note: two different drivers and two different *UAS>run* transgenes were used for these experiments (total 4 combinations) | 5B, 5D-E, 5G-H | (Mazzoni et al., 2008) |
| *run* mutant clones | *run<sup>2</sup>/(Mi ET1)MB01026* | S1A-B |  |
| *run* RNAi | *UAS>Dicer2*, *yw*; *ey>Gal4*, *GMR>Gal4/+*; *UAS>run RNAi/+* | S1D, S1F |  |
| *lz* mutants | *lz<sup>r15</sup>* | S1C |  |
| *bro* mutants | *ey>Flp*; +; *FRT80 bro/FRT80 GMR>hid*, *cL* | S1E-F | (Stowers and Schwarz, 1999) |
| *all R7s>control* | *yw*; *pm181>Gal4/+*; +/+ | S1G |  |
| *all R7s>run* | *yw*; *pm181>Gal4/+*; *UAS>run/+* | S1G |  |
| *all R7s>bro* | *yw*; *pm181>Gal4/+*; *UAS>bro/+* | S1G |  |
| *all R7s>bgb* | *yw*; *pm181>Gal4/+*; *UAS>bgb/+* | S1G |  |

### Generation of *run*^*2*^ clones

Because the *run* locus is more proximal to the centromere than *FRT19*, we created *run*^*2*^ mutant clones by radiation-induced mitotic recombination. To do so, we collected 0-24 hr old embryos that were heterozygous for *run*^*2*^ and a closely linked Minos-GFP enhancer trap ((*Mi ET1)MB01026* - which is expressed in all R7s), waited 24 hours, and then exposed them to 2 kRs of gamma-radiation. *run*^*2*^ mutant clones were indicated by the absence of GFP.

### Antibodies

Antibodies were used at the following dilutions: mouse anti-Rh3 (1:100) (gift from S. Britt, University of Colorado), rabbit anti-Rh4 (1:100) (gift from C. Zuker, Columbia University), guinea pig anti-Ss (1:500) (gift from Y.N. Jan, University of California, San Francisco), anti-Run (1:250) (gift from E. Wieschaus, Princeton University), guinea pig anti-Run (1:800) (gift from P. Gergen, Stony Brook University), sheep anti-GFP (1:500) (BioRad), chicken anti-β-gal (1:800) from Abcam (Cambridge, MA), rat anti-ELAV (1:50) (DSHB), mouse anti-Pros (1:10) (DSHB), and Alexa 488 Phalloidin (1:80) (Invitrogen). All secondary antibodies were Alexa Fluor-conjugated (1:400) and made in donkey or goat (Molecular Probes, Eugene, OR).

### Antibody staining

Adult, mid-pupal, and larval retinas were dissected as described (Hsiao et al., 2012) and fixed for 15 min with 4% formaldehyde at room temperature. Retinas were rinsed three times in PBS plus 0.3% Triton X-100 (PBX) and washed in PBX for >2 hr. Retinas were incubated with primary antibodies diluted in PBX overnight at room temperature and then rinsed three times in PBX and washed in PBX for >4 hr. Retinas were incubated with secondary antibodies diluted in PBX overnight at room temperature and then rinsed three times in PBX and washed in PBX for >2 hr. Retinas were mounted in SlowFade Gold Antifade Reagent (Invitrogen). Images were acquired using a Zeiss LSM 700 confocal microscope or a Leica SP2 microscope.

### Quantification of expression

#### Ss and Run expression in larval R7s

Expression of Ss and Run expression in PRs were manually scored in larval or pupal stages.

#### Ss and Pros expression in mid-pupal photoreceptors

Expression of Ss and Pros in PRs was assessed in mid-pupal retinas. At least 15 individual ommatidia from 5 or more mid-pupal retinas were analyzed per genotype. The number of Ss^OFF^ Pros^OFF^, Ss^ON^ Pros^ON^, Ss^OFF^ Pros^ON^, and Ss^ON^ Pros^OFF^ cells within each ommatidia were counted manually. Graphs represent average counts of each combo +-SD. Genotypes were compared to wildtype with a one-way ANOVA using a Bonferroni multiple comparison test.

#### Rh3 and Rh4 expression in adult R7s

Frequency of Rh3 (Ss^OFF^) and Rh4 (Ss^ON^) expression in R7s was scored in adults manually. Six or more retinas were scored for each genotype. 100 or more R7s were scored for each retina. Dorsal third R7s were identified by the expression of *IroC*>*LacZ*.

## Acknowledgements

We are grateful to Steve Britt, Claude Desplan, Lily Jan, Yuh-Nung Jan, Charles Zuker, J.P. Gergen, and the Bloomington and Kyoto Stock Centers for generously providing published fly stocks and antibodies. We thank members of the Johnston lab for helpful comments on the manuscript. This work was supported by NIH R01EY025598 (RJJ), NIH NRSA Genetics Training Grant 5-T32-GM007413 (ACM), and the Medical Research Foundation of Oregon (TGH).

## Figure Legends

**Figure S1.**
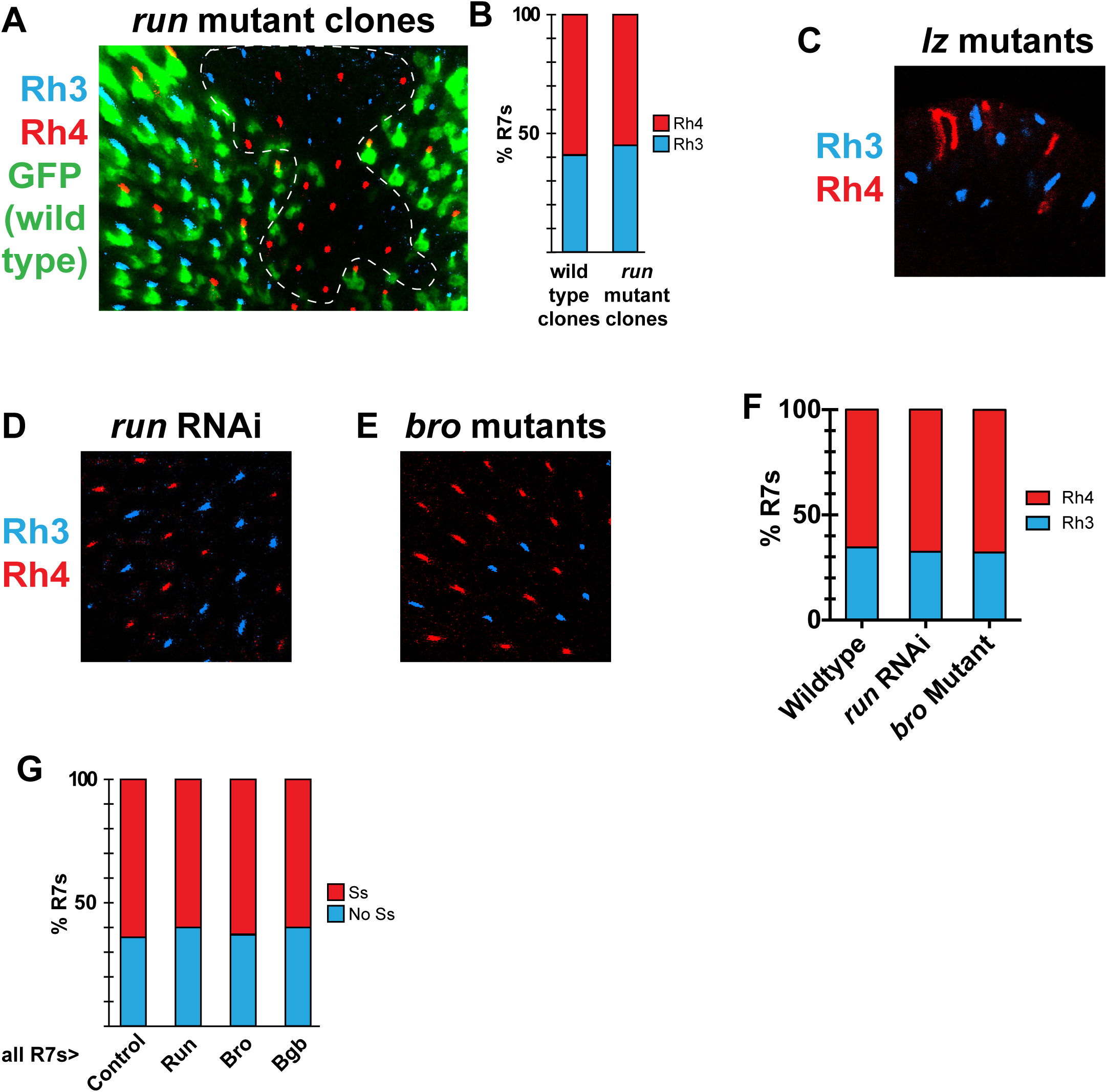
Reducing or increasing Run, Lz, Bro, or Bgb in R7s does not affect Ss, Rh3, or Rh4 expression. A. Rh3 and Rh4 expression is unaffected in *run* null mutant clones. GFP+ marks wild type clone; GFP-marks *run* null mutant clone. Dotted line indicates clone boundary. B. Quantification of A. C. Rh3 and Rh4 are stochastically expressed in the few remaining R7s in *lz* null mutants. D. Rh3 and Rh4 expression is unaffected in *run* RNAi knockdown retinas. E. Rh3 and Rh4 expression is unaffected in *bro* mutant retinas. F. Quantification of D. and E. G. Ectopic expression of Run, Bro, or Bgb in R7s does not affect Ss expression.

## References

Alqadah, A., Hsieh, Y.W., Xiong, R., Chuang, C.F., 2016. Stochastic left-right neuronal asymmetry in Caenorhabditis elegans. Philosophical transactions of the Royal Society of London. Series B, Biological sciences 371.

Anderson, C., Reiss, I., Zhou, C., Cho, A., Siddiqi, H., Mormann, B., Avelis, C.M., Deford, P., Bergland, A., Roberts, E., Taylor, J., Vasiliauskas, D., Johnston, R.J., 2017. Natural variation in stochastic photoreceptor specification and color preference in Drosophila. Elife 6.

Bell, M.L., Earl, J.B., Britt, S.G., 2007. Two types of Drosophila R7 photoreceptor cells are arranged randomly: A model for stochastic cell-fate determination. Journal of Comparative Neurology 502, 75–85.

Bessant, D.A., Payne, A.M., Mitton, K.P., Wang, Q.L., Swain, P.K., Plant, C., Bird, A.C., Zack, D.J., Swaroop, A., Bhattacharya, S.S., 1999. A mutation in NRL is associated with autosomal dominant retinitis pigmentosa. Nature genetics 21, 355–356.

Chuang, C.-F., VanHoven, M.K., Fetter, R.D., Verselis, V.K., Bargmann, C.I., 2007. An Innexin-Dependent Cell Network Establishes Left-Right Neuronal Asymmetry in C. elegans. Cell 129, 787–799.

Cook, T., Pichaud, F., Sonneville, R., Papatsenko, D., Desplan, C., 2003. Distinction between color photoreceptor cell fates is controlled by Prospero in Drosophila. Developmental cell 4, 853–864.

Daga, A., Karlovich, C.A., Dumstrei, K., Banerjee, U., 1996. Patterning of cells in the Drosophila eye by Lozenge, which shares homologous domains with AML1. Genes Dev 10, 1194–1205.

Domingos, P.M., Brown, S., Barrio, R., Ratnakumar, K., Frankfort, B.J., Mardon, G., Steller, H., Mollereau, B., 2004. Regulation of R7 and R8 differentiation by the spalt genes. Dev Biol 273, 121–133.

Duncan, D.M., Burgess, E.A., Duncan, I., 1998. Control of distal antennal identity and tarsal development inDrosophila by spineless–aristapedia, a homolog of the mammalian dioxin receptor. Genes & Development 12, 1290–1303.

Edwards, T.N., Meinertzhagen, I.A., 2009. Photoreceptor neurons find new synaptic targets when misdirected by overexpressing runt in Drosophila. The Journal of neuroscience : the official journal of the Society for Neuroscience 29, 828–841.

Furukawa, T., Morrow, E.M., Cepko, C.L., 1997. Crx, a novel otx-like homeobox gene, shows photoreceptor-specific expression and regulates photoreceptor differentiation. Cell 91, 531–541.

Hsiao, H.Y., Johnston, R.J., Jukam, D., Vasiliauskas, D., Desplan, C., Rister, J., 2012. Dissection and immunohistochemistry of larval, pupal and adult Drosophila retinas. Journal of visualized experiments : JoVE, e4347.

Hsiao, H.Y., Jukam, D., Johnston, R., Desplan, C., 2013. The neuronal transcription factor erect wing regulates specification and maintenance of Drosophila R8 photoreceptor subtypes. Dev Biol 381, 482–490.

Jacobs, G.H., Nathans, J., 2009. The evolution of Primate color vision. Sci Am 300, 56–63.

Johnston, R.J., Desplan, C., 2014. Interchromosomal Communication Coordinates Intrinsically Stochastic Expression Between Alleles. Science 343, 661–665.

Johnston, R.J., Jr., Desplan, C., 2010. Stochastic mechanisms of cell fate specification that yield random or robust outcomes. Annual review of cell and developmental biology 26, 689–719.

Johnston, R.J., Jr., Otake, Y., Sood, P., Vogt, N., Behnia, R., Vasiliauskas, D., McDonald, E., Xie, B., Koenig, S., Wolf, R., Cook, T., Gebelein, B., Kussell, E., Nakagoshi, H., Desplan, C., 2011. Interlocked feedforward loops control cell-type-specific Rhodopsin expression in the Drosophila eye. Cell 145, 956–968.

Jukam, D., Desplan, C., 2011. Binary regulation of Hippo pathway by Merlin/NF2, Kibra, Lgl, and Melted specifies and maintains postmitotic neuronal fate. Developmental cell 21, 874–887.

Jukam, D., Viets, K., Anderson, C., Zhou, C., DeFord, P., Yan, J., Cao, J., Johnston, R.J., Jr., 2016. The insulator protein BEAF-32 is required for Hippo pathway activity in the terminal differentiation of neuronal subtypes. Development 143, 2389–2397.

Jukam, D., Xie, B., Rister, J., Terrell, D., Charlton-Perkins, M., Pistillo, D., Gebelein, B., Desplan, C., Cook, T., 2013. Opposite feedbacks in the Hippo pathway for growth control and neural fate. Science 342, 1238016.

Kaminker, J.S., Canon, J., Salecker, I., Banerjee, U., 2002. Control of photoreceptor axon target choice by transcriptional repression of Runt. Nat Neurosci 5, 746–750.

Kaminker, J.S., Singh, R., Lebestky, T., Yan, H., Banerjee, U., 2001. Redundant function of Runt Domain binding partners, Big brother and Brother, during Drosophila development. Development 128, 2639–2648.

Kuhnlein, R.P., Schuh, R., 1996. Dual function of the region-specific homeotic gene spalt during Drosophila tracheal system development. Development 122, 2215–2223.

Li, L.H., Gergen, J.P., 1999. Differential interactions between Brother proteins and Runt domain proteins in the Drosophila embryo and eye. Development 126, 3313–3322.

Markenscoff-Papadimitriou, E., Allen, W.E., Colquitt, B.M., Goh, T., Murphy, K.K., Monahan, K., Mosley, C.P., Ahituv, N., Lomvardas, S., 2014. Enhancer interaction networks as a means for singular olfactory receptor expression. Cell 159, 543–557.

Mazzoni, E.O., Celik, A., Wernet, M.F., Vasiliauskas, D., Johnston, R.J., Cook, T.A., Pichaud, F., Desplan, C., 2008. Iroquois complex genes induce co-expression of rhodopsins in Drosophila. PLoS biology 6, e97.

Mears, A.J., Kondo, M., Swain, P.K., Takada, Y., Bush, R.A., Saunders, T.L., Sieving, P.A., Swaroop, A., 2001. Nrl is required for rod photoreceptor development. Nature genetics 29, 447–452.

Mikeladze-Dvali, T., Wernet, M.F., Pistillo, D., Mazzoni, E.O., Teleman, A.A., Chen, Y.W., Cohen, S., Desplan, C., 2005. The growth regulators warts/lats and melted interact in a bistable loop to specify opposite fates in Drosophila R8 photoreceptors. Cell 122, 775–787.

Miller, A.C., Seymour, H., King, C., Herman, T.G., 2008. Loss of seven-up from Drosophila R1/R6 photoreceptors reveals a stochastic fate choice that is normally biased by Notch. Development 135, 707–715.

Mlodzik, M., Hiromi, Y., Weber, U., Goodman, C.S., Rubin, G.M., 1990. The Drosophila seven-up gene, a member of the steroid receptor gene superfamily, controls photoreceptor cell fates. Cell 60, 211–224.

Mollereau, B., Dominguez, M., Webel, R., Colley, N.J., Keung, B., de Celis, J.F., Desplan, C., 2001. Two-step process for photoreceptor formation in Drosophila. Nature 412, 911–913.

Monahan, K., Lomvardas, S., 2015. Monoallelic expression of olfactory receptors. Annual review of cell and developmental biology 31, 721–740.

Morante, J., Desplan, C., Celik, A., 2007. Generating patterned arrays of photoreceptors. Curr Opin Genet Dev 17, 314–319.

Nathans, J., 1999. The evolution and physiology of human color vision: insights from molecular genetic studies of visual pigments. Neuron 24, 299–312.

Perry, M., Kinoshita, M., Saldi, G., Huo, L., Arikawa, K., Desplan, C., 2016. Molecular logic behind the three-way stochastic choices that expand butterfly colour vision. Nature 535, 280–284.

Rehemtulla, A., Warwar, R., Kumar, R., Ji, X., Zack, D.J., Swaroop, A., 1996. The basic motif-leucine zipper transcription factor Nrl can positively regulate rhodopsin gene expression. Proceedings of the National Academy of Sciences of the United States of America 93, 191–195.

Serizawa, S., Miyamichi, K., Nakatani, H., Suzuki, M., Saito, M., Yoshihara, Y., Sakano, H., 2003. Negative feedback regulation ensures the one receptor-one olfactory neuron rule in mouse. Science 302, 2088–2094.

Smallwood, P.M., Wang, Y., Nathans, J., 2002a. Role of a locus control region in the mutually exclusive expression of human red and green cone pigment genes. Proceedings of the National Academy of Sciences of the United States of America 99, 1008–1011.

Smallwood, P.M., Wang, Y., Nathans, J., 2002b. Role of a locus control region in the mutually exclusive expression of human red and green cone pigment genes. Proceedings of the National Academy of Sciences 99, 1008–1011.

Stowers, R.S., Schwarz, T.L., 1999. A genetic method for generating Drosophila eyes composed exclusively of mitotic clones of a single genotype. Genetics 152, 1631–1639.

Thanawala, Shivani U., Rister, J., Goldberg, Gregory W., Zuskov, A., Olesnicky, Eugenia C., Flowers, Jonathan M., Jukam, D., Purugganan, Michael D., Gavis, Elizabeth R., Desplan, C., Johnston Jr, Robert J., 2013. Regional Modulation of a Stochastically Expressed Factor Determines Photoreceptor Subtypes in the Drosophila Retina. Developmental cell 25, 93–105.

Troemel, E.R., Sagasti, A., Bargmann, C.I., 1999. Lateral Signaling Mediated by Axon Contact and Calcium Entry Regulates Asymmetric Odorant Receptor Expression in C. elegans. Cell 99, 387–398.

Urban, E.A., Johnston, R.J., Jr., 2018. Buffering and Amplifying Transcriptional Noise During Cell Fate Specification. Front Genet 9, 591.

Viets, K., Eldred, K.C., Johnston, R.J., Jr., 2016. Mechanisms of Photoreceptor Patterning in Vertebrates and Invertebrates. Trends Genet 32, 638–659.

Wernet, M.F., Mazzoni, E.O., Çelik, A., Duncan, D.M., Duncan, I., Desplan, C., 2006. Stochastic spineless expression creates the retinal mosaic for colour vision. Nature 440, 174–180.

Wernet, M.F., Perry, M.W., Desplan, C., 2015. The evolutionary diversity of insect retinal mosaics: common design principles and emerging molecular logic. Trends Genet 31, 316–328.

Xie, B., Charlton-Perkins, M., McDonald, E., Gebelein, B., Cook, T., 2007. Senseless functions as a molecular switch for color photoreceptor differentiation in Drosophila. Development 134, 4243–4253.

Yan, J., Anderson, C., Viets, K., Tran, S., Goldberg, G., Small, S., Johnston, R.J., Jr., 2017. Regulatory logic driving stable levels of defective proventriculus expression during terminal photoreceptor specification in flies. Development 144, 844–855.

